# Stress-relief function of migrasomal autophagosome formed by inhibition of autophagosome/lysosome fusion

**DOI:** 10.1101/2022.06.08.495398

**Authors:** Seon Yong Lee, Sang-Hun Choi, Hee-Sung Ahn, Young-Gyu Ko, Kyunggon Kim, Sung Wook Chi, Hyunggee Kim

## Abstract

Glioblastoma (GBM) is more difficult to treat than other intractable adult tumors. Here, we describe the composition and function of migrasomes generated along with GBM cell migration. Proteomic analysis revealed that LC3B-positive autophagosomes were abundant in the migrasomes of GBM cells. An increased number of migrasomes was observed following treatment with chloroquine (CQ) or inhibition of the expression of *STX17* and *SNAP29*, which are involved in autophagosome/lysosome fusion. Although *ATG7* ablation, which is involved in LC3B lipidation, did not suppress migrasome formation, it was confirmed that migrasome formation could be diminished by blocking the alternative autophagy pathway through double knockout of *ATG7*/*BECN1*. Furthermore, depletion of *ITGA5* or *TSPAN4* did not relieve endoplasmic reticulum (ER) stress in cells, resulting in cell death. Taken together, our study suggests that increasing the number of autophagosomes, through inhibition of autophagosome/lysosome fusion, generates migrasomes that have the capacity to alleviate cellular stress.

**Summary statement:** This study demonstrates that glioblastoma cells contain autophagosomes within their migrasomes. Under stress conditions, the formation of migrasomes serves as a stress-relief mechanism to alleviate cell death.

## Introduction

Glioblastoma (GBM) is a refractory tumor with an incidence of approximately ∼16% among all adult primary brain tumors (Ostrom et al., 2013). The median survival is only 12-14 months, and the only standard treatment is a combination of temozolomide and ionizing radiation therapy (Bohn et al., 2018; Stupp et al., 2009; Stupp et al., 2005). Recurrence of GBM, with a probability of up to 90%, also contributes to poor prognosis for patients with GBM (Weller et al., 2013). Therefore, there have been many studies and efforts to treat GBM (Bagley et al., 2018; Cloughesy et al., 2019; Lombardi et al., 2019).

In 2015, the Li Yu group discovered a multivesicular body-like structure on the trailing edge of a migrating cell (Ma et al., 2015). They named it the “Migrasomes.” Migrasomes are generated during cell movement and are randomly distributed inside thin cellular structures called retraction fibers (RFs). These structures have been found in various types of cells, including normal rat kidney cells, and recently, it has been suggested that their cell membranes are composed of tetraspanin- and cholesterol-enriched microdomains (TEMAs) (Huang et al., 2019; Ma et al., 2015). The cell membrane composed of TEMAs is guided by the stiffening phenomenon of the membrane caused by adhesion to the substrate on the bottom surface of the cultured cells, and this physical interaction creates a unique cell membrane structure (Huang et al., 2019). This newly discovered cell membrane structure, which was thought to be reproduced only under artificial experimental conditions, reached a milestone when its formation and function were suggested in the zebrafish gastrulation phase (Jiang et al., 2019). That is, the migrasome supplying CXCL12 chemokine is generated in mesodermal and endodermal cells, and it was shown that chemotaxis by CXCL12 induces normal migration of dorsal forerunner cells (DFCs). As such, it has been established that migrasomes play an important role in the development of organisms, and the investigation of their function in vivo will enhance interest regarding their role in other fields of research. Recently, it has been demonstrated in neutrophils that mild mitochondrial stress leads to the incorporation of damaged mitochondria into the migrasomes, in a process called “mitocytosis.” (Jiao et al., 2021) Simultaneously, interest in RF, which was only considered a pathway to construct the migrasome, is being highlighted (Wang et al., 2022).

During nutrient starvation or proteostatic stress, to survive under harsh conditions, cells restrict their energy use and produce essential components for maintaining cell viability using the autophagic pathway (Lim et al., 2015; Russell et al., 2014). During autophagy, cells produce double-membrane structures, namely phagophores, usually from the endoplasmic reticulum (ER), and multiple autophagy-related (ATG) genes are involved in this process (Juhasz and Neufeld, 2006). It is known that about 20 ATG proteins are involved in the canonical autophagy pathway (Dikic and Elazar, 2018). The core ATG proteins include the ULK1/2 kinase complex, ATG9A-associated vesicle trafficking system, and the PI3KC3 (Vps34 in yeast) complex (Klionsky et al., 2010). These protein complexes play an essential role in phagophore nucleation during autophagy initiation. LC3 (*MAP1LC3A*, *MAP1LC3B*, and *MAP1LC3C*) is involved in the phagophore maturation stage (Dikic and Elazar, 2018; Klionsky et al., 2010). In yeast, it is an ubiquitin-like protein known as Atg8 (Farré and Subramani, 2016). The E1-like enzyme ATG7 adenylates ATG12 and is involved in the formation of ATG5-ATG12 conjugates, whose formation is mediated by the E2-like enzyme ATG10 (Dikic and Elazar, 2018; Rubinsztein et al., 2012). In addition, ATG7 plays a role in the phosphatidylethanolamine conjugation of LC3 and recruit LC3 to the phagophore, which is important for phagophore elongation (Weidberg et al., 2010). Upon formation, autophagosomes fuse with the lysosome with the assistance of the SNARE protein complex called STX17-SNAP29-VAMP8, resulting in lysosomal degradation of substances within autophagosomes (Itakura et al., 2012; Shen et al., 2021).

Autophagy plays various roles depending on the tumor type, stage of cancer progression, and the genomic status of each tumor (Li et al., 2020; Yun and Lee, 2018). Autophagy also acts as the quality control process for proteins and organelles, helping cancer cells evade stress response and suppressing inflammatory responses of immune cells in tumor lesion (Kroemer et al., 2010; Ogata et al., 2006). In contrast, in mouse models in which ATG proteins (ATG7 and Beclin-1) are suppressed, tumorigenic potential is promoted, and spontaneous tumors can be formed (Santanam et al., 2016; Yue et al., 2003). In particular, Beclin-1 has been shown to function as a haploid-insufficient tumor suppressor in hepatocellular carcinoma, prostate cancers, and breast cancers (Liang et al., 2018; Liang et al., 1999; Liu et al., 2013; Wijshake et al., 2021). Accumulated sequestosome 1 (SQSTM1, i.e., p62) protein is also found in gastrointestinal carcinoma and prostate cancer, and the accumulated p62 protein is used as a marker for malignant tumors (Jiang et al., 2020; Nakayama et al., 2017; Zhu et al., 2018).

As described above, the autophagy pathway also acts as a stress-relief mechanism in cancer cells exposed to various types of stressors (Huang et al., 2016). Cancer cells are exposed to environmental conditions such as chronic nutrient deficiency, hypoxia, and low pH due to abnormal blood vessel formation (Fukumura and Jain, 2007; Lugano et al., 2020). In these environments, autophagy activation may play a crucial role in cancer progression and development. Autophagy is also known to be activated even after GBM treatment (Knizhnik et al., 2013; Wang et al., 2018). Several studies have reported that the autophagy pathway suppresses GBM cell death (Hori et al., 2015; Katayama et al., 2007). Treatment options for patients with GBM, such as arsenic trioxide (As_2_O_3_), temozolomide, rapamycin, and tamoxifen, promote autophagy to alleviate GBM cell death (Graham et al., 2016; Kanzawa et al., 2004; Kanzawa et al., 2005; Takeuchi et al., 2005).

The functions of RF and the migrasome (R&M) have not been completely elucidated, and in particular, their role in cancer biology has not been investigated. In this study, we investigated the relationship between autophagosome and R&M formation and identified that R&M have a stress-relief function in brain tumor cells upon ER stress condition.

## Result

### Identification of cellular organelles enriched in GBM cell-derived R&Ms

RFs and migrasomes of GBM cells have been described in our previous study and by the Li Yu group (Lee et al., 2021; Ma et al., 2015). We anticipated that vigorously migrating GBM cells would generate migrasomes behind their trailing edge. We visualized RFs previously through CD9, a tetraspanin protein (Huang et al., 2019). CD9 overexpression did not increase the number of migrasomes as in case of *TSPAN4* overexpression. We found that some cells were capable of generating migrasomes, even though other cells exhibited fewer migrasomes and only produced RFs (Fig. S1A). In 2021, Wang et al. investigated the presence of damaged mitochondria within neutrophil migrasomes, both in vitro and in vivo (Jiao et al., 2021). Thus, we were intrigued as to which cellular organelles are abundantly located in migrasomes. For performing proteomic analysis, we purified crude migrasomes including RFs (i.e., R&M) along with extracellular vesicles (EVs) using serial centrifugation (Fig. 1A; Fig. S1B-D). We then analyzed the cellular materials to distinguish them according to their size. The diameter of R&Ms were about 1.7-fold larger than that of EVs (EV: 177.7 ± 4.1 nm, R&M: 307.7 ± 6.0 nm), suggesting that crude migrasomes are large enough to distinguish them from EVs (Fig. 1B). Moreover, we further analyzed the relative abundance of proteins using principal component analysis based on liquid chromatography-high resolution mass spectrometry (LC-HRMS) (Fig. 1C,D). Technical replicates of each sample revealed a separate distribution in the analysis. We then examined the specific proteins enriched in each sample by calculating the relative abundance of proteins within the samples after adjusting the raw abundance value of the proteins. We defined the proteins enriched in each sample as those with a relative abundance value above the first quartile, “Q1” (Fig. 1D). By using combined gene sets from “common Q1” and “R&M-specific Q1”, we performed enrichment analysis using Enrichr, which is powered by Appyter (Chen et al., 2013; Kuleshov et al., 2016). Consequently, enrichment analysis using GO-Cellular Component v2021 revealed enriched cellular organelles of combined gene sets to dimensionality-reduced uniform manifold approximation and projection (UMAP) (Fig. 1E). Among the diverse cellular organelles, autophagosome/lysosome, ER/vesicle, and cytoskeleton were the major organelles in the R&M samples.

**Fig. 1.**
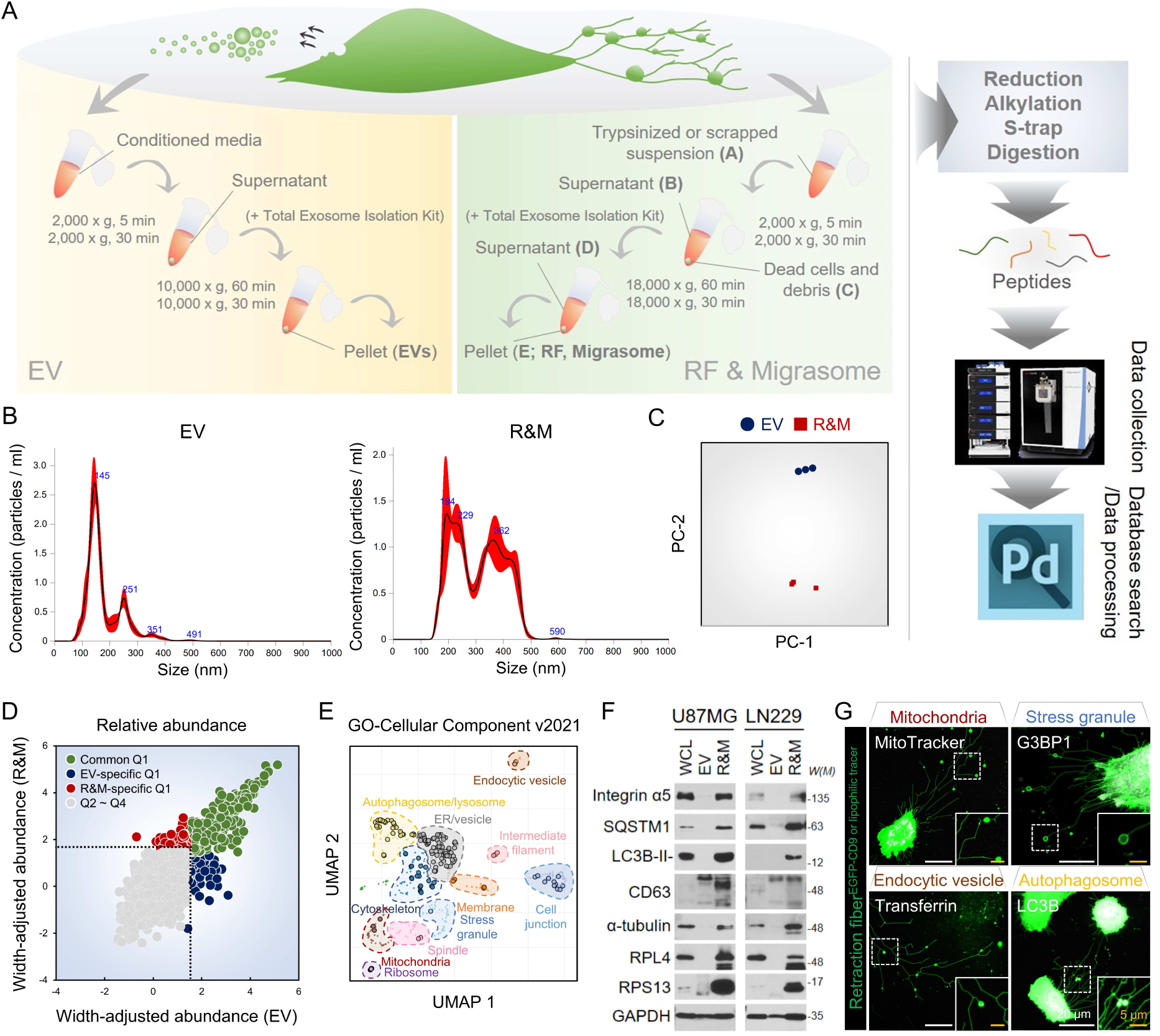
Identifying the presence of the autophagosomes within the retraction fiber & migrasome (R&M) of glioblastoma cells. (A) Purification procedures of tumor-derived extracellular vesicle (EV) and R&M. Both samples are purified and further analyzed using liquid chromatography-high resolution mass spectrometry (LC-HRMS, Orbitrap Exploris 480). Samples were divided into three-technically replicated sub-samples. For database searching and processing, we searched mapped protein sequences against SwissProt database (release v2019_06) and Proteome discoverer v2.2. (B) Nano particle tracking analysis used for quantifying and qualifying both EV and R&M. Three biological replicates were used for analysis. (C) Principal component analysis (PCA) shows that EVs and R&Ms have proteins with a distinctly different composition. PCA was performed using relative protein abundance values. The raw values of each protein abundance were converted to log2 values. Then, the amount of protein was corrected using the width adjustment method. (D) Based on the width-adjusted relative protein abundance values, the protein present in the first quartile or higher in each sample was specified as “Q1”. The remaining proteins, belonging to “Q2 – Q4”, were excluded from the predominantly present proteins in each sample. (E) A scatter plot visualizing the result of gene ontology enrichment analysis (GO-Cellular Component v2021) for common Q1 and R&M-specific Q1 combined gene set. Plot was obtained from Enrichment Analysis Visualization Appyter v0.2.5 and it organized similar gene ontology gene sets into clusters using first two UMAP dimensions. We manually designated each cluster by follows: mitochondria, ribosome, stress granule, membrane, cell junction, intermediate filament, endocytic vesicle, ER/vesicle, cytoskeleton, spindle, autophagosome/lysosome. (F) Western blotting of integrin α5, SQSTM1, LC3B, CD63, α-tubulin, RPL4, RPS13, and GAPDH proteins of whole cell lysate (WCL), EV, and R&M in U87MG and LN229 cells. *W(M)*, molecular weight. (G) Live-cell imaging for visualizing marker proteins of major cellular organelles derived from the result of Fig. 1E. White, each marker of cellular organelles; MitoTracker Deep Red for visualizing mitochondria, tdTomato-G3BP1 for visualizing stress granules, EGFP-LC3B for visualizing autophagosomes, and transferrin 488-conjugate for visualizing endocytic vesicles. Green, EGFP-CD9 or DiO/DiI lipophilic tracer. Scale bars (white), 20 μm. Scale bars (yellow), 5 μm.

Next, we investigated whether the R&M-enriched cellular organelles were truly located within R&M. Thus, we confirmed the expression of representative marker proteins using western blot analysis and live-cell fluorescence imaging (Fig. 1F,G). Integrin α5, α-tubulin, CD63, and GAPDH are known marker proteins of migrasomes and exosomes (Huang et al., 2019; Lee et al., 2021; Zhu et al., 2021). Moreover, G3BP1, a major protein for nucleating stress granules (Panas et al., 2016), and mitochondria, which are known to be transported through retraction fibers in an anterograde manner during mitochondrial damage (Jiao et al., 2021), were not observed in steady-state of cells. Transferrin, located in endocytic vesicles (van Renswoude et al., 1982), has been detected in a few migrasomes. Interestingly, both LC3B-II, the lipidated form of LC3B (*MAP1LC3B* for gene name), and autophagosome bridging protein, SQSTM1, were highly expressed in R&M samples from both U87MG and LN229 cells. EGFP-LC3B was also strongly observed in live-cell imaging. Purified R&M samples were also prepared for detection using transmission electron microscopy. Some polysome-like structures, and lipid droplets, and autophagosome-like structures were observed (Fig. S1E). Together, we analyzed enriched cellular organelles within the R&M of GBM cells and observed high expression of LC3B and SQSTM1 proteins in R&M samples and intact migrasomes.

### Inhibition of autophagosome/lysosome fusion promotes the formation of R&Ms of GBM cell

Next, we investigated whether the LC3B-II proteins found in the R&M samples of GBM cells were derived from the cellular autophagy process. As the phosphatidylethanolamine-conjugated LC3B-II proteins are accumulated in the circumstance of autophagic flux inhibition (Mauthe et al., 2018), we examined the formation of R&M in the presence of chloroquine (CQ) or bafilomycin A1 (BafA1), which disrupt autophagic degradation of cargos within autolysosomes under normal culture conditions. Both CQ and BafA1 treated samples exhibited increased levels of LC3B-II (Fig. S2A), as previously reported (Mauthe et al., 2018). However, only CQ-treated LN229 cells generated more R&Ms compared to vehicle-treated cells (Fig. 2A-C). BafA1 is a vacuolar H^+^-ATPase inhibitor that inhibits the entry of H^+^ into lysosomes, thereby disrupting their function. CQ inhibits autophagosome/lysosome fusion; thus, pH-neutral LC3B-positive autophagosomes are concentrated in the native endomembrane system (Mauthe et al., 2018). Therefore, we expressed a tandem-fluorescent LC3B construct (Klionsky et al., 2021) in LN229 cells to determine whether complete autophagosomes, which do not fuse with lysosomes, are observed within the R&M. Consequently, we observed increased number of LC3B-positive vesicles in both the cell body and migrasomes in the CQ-treated group (Fig. 2D,E). These LC3B-positive vesicles were also dual fluorescent-positive, indicating pH-neutral autophagosomes existed within migrasomes. Next, we hypothesized that autophagosomes at the stage of phagophore maturation were observed within migrasomes (Fig. 2F). During phagophore maturation, the E1-like enzyme ATG7 mediates the conjugation of ATG5 and ATG12 to render phosphatidylethanolamine to LC3B-I proteins via complex formation with ATG16L (Klionsky et al., 2010; Rubinsztein et al., 2012). Thus, we performed a proximity ligation assay for LC3B and ATG5 (Fig. 2G). LC3B and ATG5 interact with each other; thus, autophagosomes, but not autolysosomes, are identified as sequestrated cellular vesicles during CQ treatment.

**Fig. 2.**
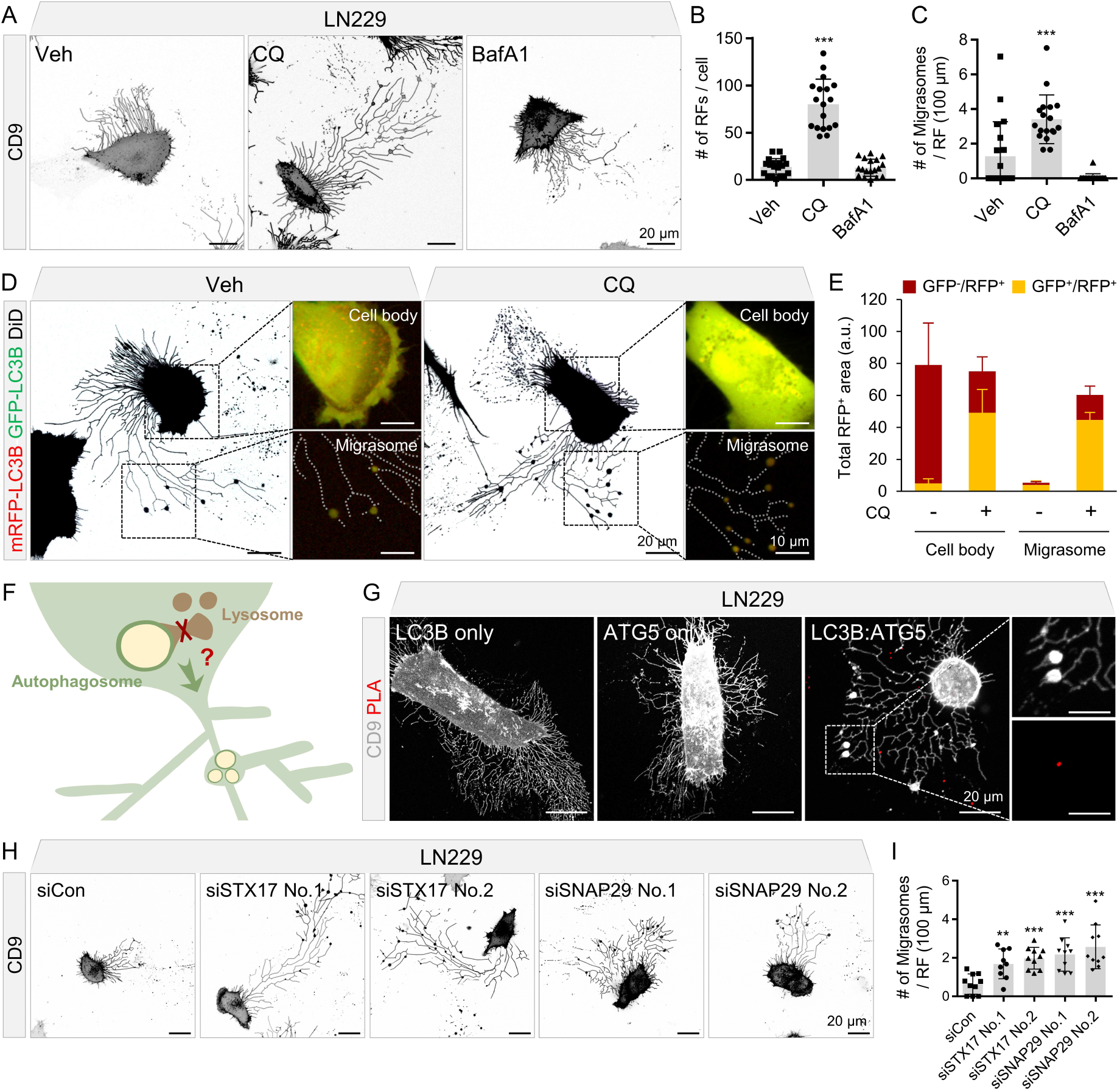
Inhibition of autophagosome/lysosome fusion induces retraction fiber and migrasome (R&M) formation. (A) Live-cell imaging for observing R&Ms formed in LN229 cells. Cells were treated with chloroquine (CQ; 50 μM, 12 h) or bafilomycin A1 (BafA1; 50 nM, 12 h). Gray, EGFP-CD9. Scale bars, 20 μm. (B) Quantification of the number of retraction fibers (RFs) per a cell. Image analyses were performed using results from Fig. 2A. *** indicates p < 0.001. Data are expressed as mean ± SEM. The unpaired nonparametric Mann-Whitney U test was used to analyze the statistical significance between each group (n = 19 for vehicle-treated condition, n = 18 for CQ- or BafA1-treated condition). The figure is representative of three-biological replicates with similar results. (C) Quantification of the number of migrasomes per RF (100 μm). Image analyses were performed using the results from Fig. 2A. *** indicates p < 0.001. Data are expressed as mean ± SEM. The unpaired nonparametric Mann-Whitney U test was used to analyze the statistical significance between each group. (D) Tandem-fluorescent LC3B was expressed in LN229 cell. GFP^-^/RFP^+^ vesicles represent autolysosome and GFP^+^/RFP^+^ vesicles represent autophagosome. Fluorescent signal observed in both cell body and migrasome were quantified respectively. Cells were treated with CQ (50 μM, 12 h). Red/Green, mRFP-EGFP-LC3B. Black, DiD lipophilic tracer. Scale bars (black), 20 μm. Scale bars (white), 10 μm. (E) Quantification of Fig. 2D. Total RFP^+^ vesicle area was quantified in each condition. The figure is representative of three-biological replicates with similar results. (F) Hypothetic graphical scheme of the relationship between autophagosome/lysosome fusion and R&M formation. (G) In situ proximity ligation assay (PLA) was performed to examine the interaction between LC3B and ATG5 in migrasomes of LN229. Cells were treated with CQ (50 μM, 12 h). Red, PLA signal. Gray, EGFP-CD9. Scale bars, 20 μm. Scale bars in cropped panels, 10 μm. (H) Live-cell imaging for observing retraction fiber (RF) & migrasome formed in LN229 cells expressing EGFP-CD9 after transfection of *STX17* or *SNAP29* siRNAs. Gray, EGFP-CD9. Scale bars, 20 μm. (I) Quantification of the number of migrasomes per RF (100 μm). Image analyses were performed using the results from Fig. 2H. ** indicates p < 0.01. *** indicates p < 0.001. Data are expressed as mean ± SEM. The unpaired nonparametric Mann-Whitney U test was used to analyze the statistical significance between each group (n = 10).

It is known that CQ abrogates autophagosome/lysosome fusion in association with the separation of STX17-positive vesicles from LAMP-2-positive lysosomes, which means that complete autophagosomes cannot undergo vesicular fusion with lysosomes in the presence of CQ (Mauthe et al., 2018). Autophagosome/lysosome fusion is mediated by the STX17-SNAP29-VAMP8 complex (Shen et al., 2021). STX17 is a SNARE protein that binds to the complete autophagosome and recruits SNAP29, which mediate complex formation of SNARE proteins with both autophagosomes (STX17) and lysosomes (VAMP8) (Itakura et al., 2012; Shen et al., 2021). Thus, we depleted *STX17* and *SNAP29* expression in LN229 cells using small interfering RNA (siRNA) to investigate whether the inhibition of autophagosome/lysosome fusion by ablation of SNARE proteins induced the formation of R&M. As a result, the R&Ms of LN229 cells were increased in both *STX17* and *SNAP29* ablated condition (Fig. 2H,I; Fig. S2B,C). Altogether, we identified that increased R&M after treatment with CQ is associated with inhibition of the autophagosome/lysosome fusion process, suggesting that autophagosomes that cannot be fuse with lysosomes are present within migrasomes.

### Ribosomes are abundant cellular cargo proteins in R&Ms

Tumor cells require vigorous amounts of building blocks to survive under stressful conditions, such as nutrient depletion and hypoxic conditions (Lim et al., 2015; Russell et al., 2014). Thus, these cells activate the autophagy pathway to enhance cell viability. Several macromolecules and organelles are involved in autophagic degradation (Dikic and Elazar, 2018; Juhasz and Neufeld, 2006; Klionsky et al., 2010). We investigated cargo proteins that were abundantly present in R&M samples. Using “R&M-specific Q1 (Fig. 1D)” as set of genes for identifying major cargo proteins within R&M, we found that cytosolic ribosomes, including both large and small subunits of ribosomal proteins, were specifically present in the R&M samples. These ribosomal subunit genes were also grouped using GeneMANIA analysis (Fig. S3A). We performed ClueGO analysis which shows relevant gene ontology terms that interact with edges and indicated statistical significance with colored nodes (Fig. S3B). Several ribosomal subunit genes were more highly expressed in the R&M samples than in the EV samples (Fig. 3A). Thus, we cloned both the large and small ribosomal subunit genes (*RPLP2* and *RPS13*) to confirm whether they were present within R&M. We expressed tdTomato-RPLP2 or RPS13 genes in EGFP-CD9 expressing GBM cells. We observed that both ribosomal proteins were located within migrasomes (Fig. 3B; Fig. S3C). Ribosomes constitute the translational machinery located in the rough ER and play essential roles in protein synthesis (Pestova et al., 2001). During eukaryotic translation initiation, 40s small and 60s large ribosomal subunits are sequentially assembled to the 5’-UTR of mRNA by translation initiation factors to generate 80s ribosomes (Pestova et al., 2001). However, in the context of proteotoxic stress, such as low-dose arsenic stress, transient ER stress occurs concomitantly with the inhibition of autophagy, and ribophagic flux is promoted in an ATG5-dependent manner (An and Harper, 2018; Dodson et al., 2018a). Thus, we hypothesized that arsenic stress evokes ribosome stress combined with ER stress and inhibits total autophagic flux, like in case of CQ treatment. We treated LN229 cells expressing EGFP-CD9 and tdTomato-RPS13 with CQ, sodium arsenite (NaAsO_2_; AS), and a combination of both. Both CQ and AS upregulated p-eIF2a levels, indicating that ER stress was induced, and CQ and AS treatment had the same inhibitory effects on autophagy flux; that is, both conditions inhibited the autophagy pathway and prevented autophagosomes from fusing with lysosomes (Fig. 3C). It has been reported that arsenic-induced autophagy inhibition is mediated by increased O-GlcNAcylation of the SNAP29 protein (Dodson et al., 2018b). Thus, in this case, AS may inhibit autophagosome/lysosome fusion via the same mechanism as CQ. As we observed that cells in the AS treatment condition exhibited an increased number and size of migrasomes, including the number of RFs (Fig. 3D-F). We also detected dual-fluorescent LC3B-positive autophagosomes within migrasomes (Fig. S2D). Together, we determined that ribosomal subunit proteins were predominantly enriched within the R&Ms of GBM cells. Proteotoxic stress, such as that caused by AS, induces ER stress with an inhibitory effect on autophagy, suggesting that ribosomes are degraded under stressful conditions However, these proteins cannot be completely degraded and left in autophagosomes (i.e., located within migrasomes) upon exposure to CQ or AS due to their inhibitory effect on autophagy pathway.

**Fig. 3.**
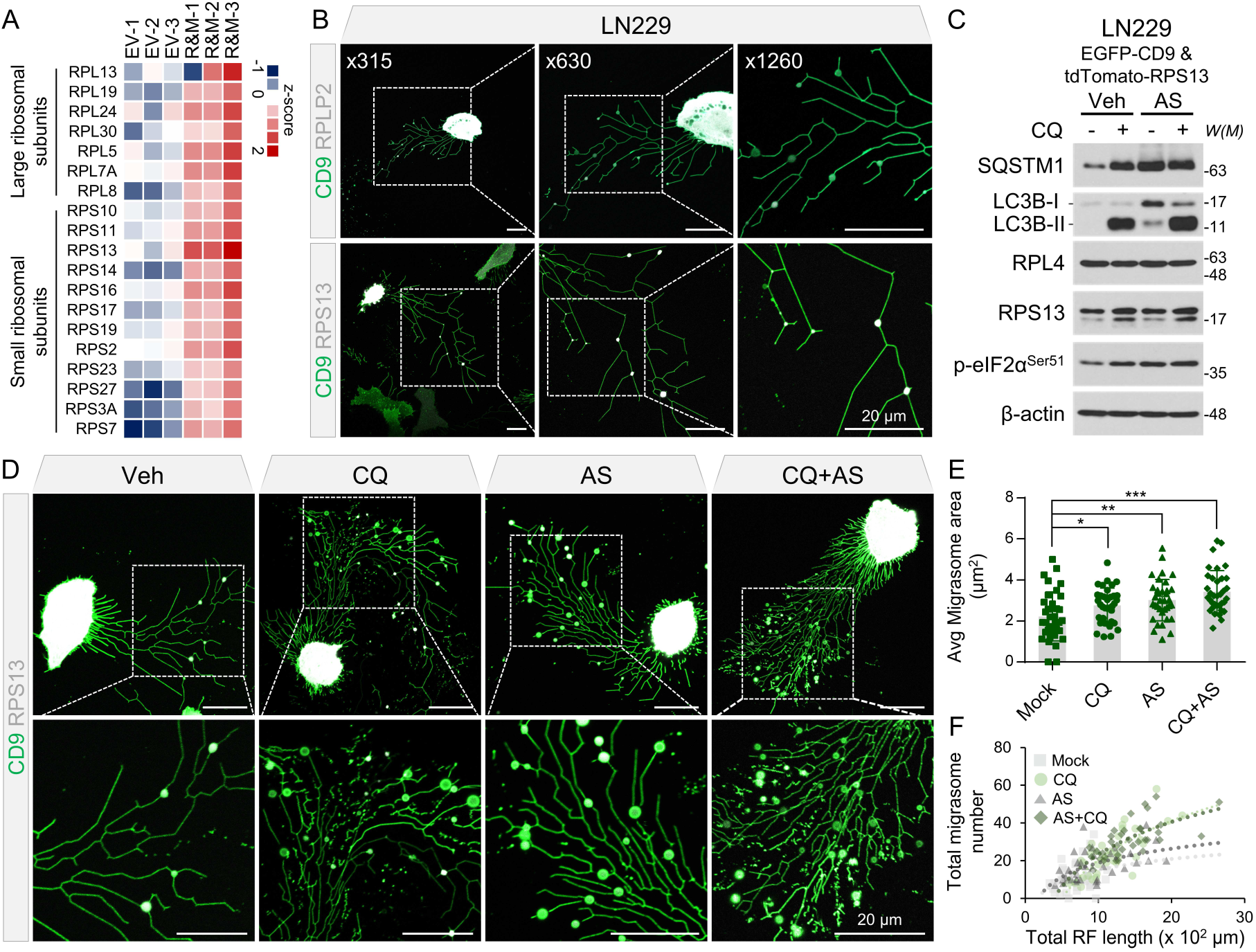
Ribosomes are abundant cargo proteins present in retraction fiber & migrasome (R&M). (A) Heatmap represents highly enriched ribosomal subunits proteins in R&M portion. Relative abundance values of both extracellular vesicle (EV) and R&M samples were standardized to z-score. Both large and small subunits of ribosomal proteins are relatively enriched in R&M. (B) Live-cell imaging of LN229 cells for visualizing the ribosomal proteins (RPLP2 and RPS13) present in migrasome. Green, EGFP-CD9. White, ribosomal proteins (tdTomato-RPLP2 or tdTomato-RPS13). Scale bars, 20 μm. (C) Western blotting of SQSTM1, LC3B, RPL4, RPS13, p-eIF2α (Ser51), and β-actin proteins after treatment of chloroquine (CQ; 50 μM, 12 h) or NaAsO_2_ (AS; 10 μM, 12 h) in LN229 (EGFP-CD9 & tdTomato-RPS13) cells. *W(M)*, molecular weight. (D) Live-cell imaging for quantifying R&M formation in CQ (50 μM, 12 h) or AS (10 μM, 12 h) treatment condition. Green, EGFP-CD9. White, tdTomato-RPS13. Scale bars, 20 μm. (E) Quantification of average migrasome area (μm^2^) in data from Fig. 3D. LN229 (EGFP-CD9 & tdTomato-RPS13) cell was used for observation and image analyses. * indicates p < 0.05; ** indicates p < 0.01; *** indicates p < 0.001. Data are expressed as mean ± SEM. The unpaired nonparametric Mann-Whitney U test was used to analyze the statistical significance between each group. (F) Quantification of total migrasome number and total RF length (x 10^2^ μm) in data from Fig. 3D.

### Ablation of ATG genes disrupts R&M formation in GBM cells

Inhibition of autophagic flux concentrated the level of LC3B-positive autophagosomes and sequentially generated more R&Ms in GBM cells (Fig. 2; Fig. 3). We then focused on how ATG genes are involved in R&M formation. First, we knocked out some of core ATG genes (*ATG5*, *ATG7*, and *BECN1*) in LN229 cells to disrupt autophagosome maturation and phagophore nucleation (Fig. S4A). We established cell lines expressing each guide RNA for CRISPR knockout and treated each cell with CQ to inhibit autophagic flux and induce R&M formation. However, we did not observe any noticeable changes in the number of R&Ms, even though we confirmed elevated levels of SQSTM1 and inhibition of LC3B-II levels in *ATG5* or *ATG7* knockout cell lines due to dysfunction of LC3B lipidation (Fig. 4A-D). Beclin-1 is associated with phagophore nucleation in the form of a class III PI3K (PI3KC3-Beclin-1) complex, and this autophagic process is mediated by a PI3P-binding complex-independent manner, indicating that alternative membrane nucleation complemented the autophagy pathway (Li et al., 2008). Thus, we suppressed both core membrane nucleation complex by knocking out both *ATG7* and *BECN1* genes in LN229 cells to investigate R&M formation (Fig. 4E,F). Intriguingly, we observed a lower number of migrasomes in *ATG7*/*BECN1* double knockout cells than in control or *ATG7* knockout cells (Fig. 4G-I). Taken together, we investigated whether ATG genes associated with phagophore initiation and maturation were involved in R&M formation in GBM cells. We found that the simultaneous disruption of phagophore nucleation and maturation diminished the formation of R&M and enhanced stress response.

**Fig. 4.**
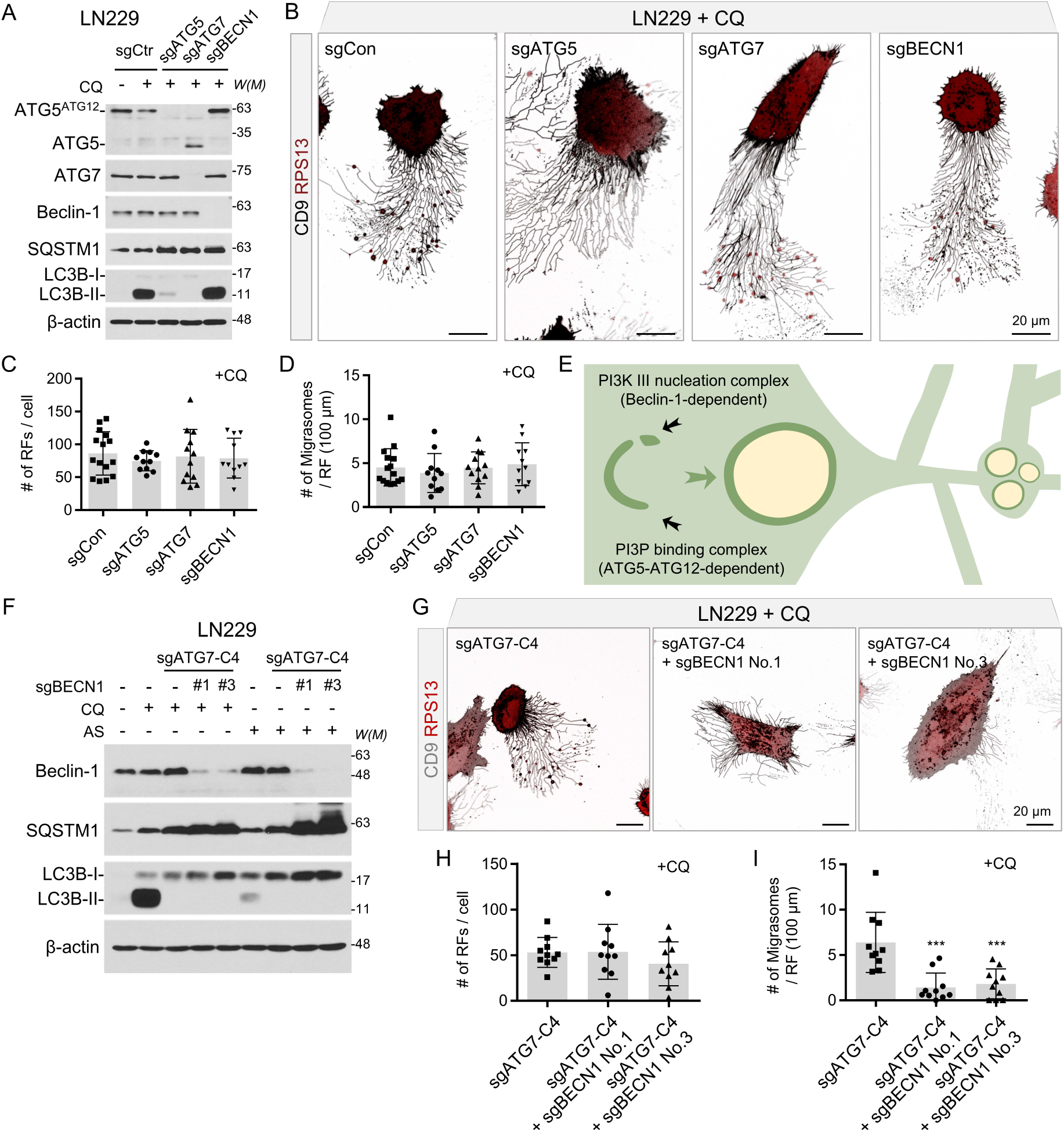
Ablation of autophagy-related (ATG) genes disrupts retraction fiber & migrasome (R&M) formation. (A) Knockout of ATG genes (*ATG5*, *ATG7*, and *BECN1*) in LN229 cell. Western blotting of ATG5 (ATG5-ATG12 conjugation form), ATG7, Beclin-1, SQSTM1, LC3B, and β-actin proteins in each autophagy related gene-sgRNA (each sgRNA using No.2 sgRNA in Fig. S4A) transduced LN229 cells. Cells were treated with chloroquine (CQ; 50 μM, 12 h). *W(M)*, molecular weight. (B) Live-cell imaging of ATG gene (*ATG5*, *ATG7*, and *BECN1*) knockout LN229 cells. Cells were treated with CQ (50 μM, 12 h). Gray, EGFP-CD9. Red, tdTomato-RPS13. Scale bars, 20 μm. (C) Quantification of the number of RFs per a cell. Image analyses were performed using results from Fig. 4B. Data are expressed as mean ± SEM. The unpaired nonparametric Mann-Whitney U test was used to analyze the statistical significance between each group (n = 10). (D) Quantification of the number of migrasomes per RF (100 μm). Image analyses were performed using results from Fig. 4B. Data are expressed as mean ± SEM. The unpaired nonparametric Mann-Whitney U test was used to analyze the statistical significance between each group (n = 10). (E) Graphical hypothesis of migrasome formation and its association with core ATG proteins. (F) Western blotting of control knockout, *ATG7* knockout single cell clone (Clone #4), and *ATG7*/*BECN1* double-knockout LN229 cells in CQ (50 μM, 12 h) or NaAsO_2_ (AS; 10 μM, 12 h) treatment condition. The protein levels of ATG7, Beclin-1, SQSTM1, LC3B, and β-actin proteins were examined. (*BECN1* sgRNA using No.1 or No.3 sgRNA in Fig. S4A). *W(M)*, molecular weight. (G) Live-cell imaging of control knockout, *ATG7* knockout single cell clone (Clone #4), and *ATG7*/*BECN1* double-knockout cells in treatment with CQ (50 μM, 12 h). Gray, EGFP-CD9. Red, tdTomato-RPS13. Scale bars, 20 μm. (H) Quantification of the number of RFs per a cell. Image analyses were performed using results from Fig. 4G. Data are expressed as mean ± SEM. The unpaired nonparametric Mann-Whitney U test was used to analyze the statistical significance between each group (n = 10). (I) Quantification of the number of migrasomes per RF (100 μm). Image analyses were performed using results from Fig. 4G. *** indicates p < 0.001. Data are expressed as mean ± SEM. The unpaired nonparametric Mann-Whitney U test was used to analyze the statistical significance between each group (n = 10).

### Inhibition of R&M formation induces cell death through out-of-control of ER stress

Previously, it has been reported that the autophagy pathway is associated with ER stress (Qi and Chen, 2019). Moreover, inhibition of autophagosome/lysosome fusion induced ER stress concomitant with R&M formation (Fig. 3C). Next, we disturbed R&M formation by decreasing the levels of *ITGA5* and *TSPAN4* (Huang et al., 2019; Lee et al., 2021; Zhu et al., 2021), regulators of R&M formation to investigate the function of R&M in ER stress condition. By depleting the levels of both *ITGA5* and *TSPAN4* using siRNA, we observed a lower number of R&M in U87MG cells than in control cells (Fig. 5A-F). Furthermore, we examined whether reduced R&M formation renders cells vulnerable to ER stress. Thus, we analyzed cellular phenotypes after stimulation with CQ or AS to induce ER stress concomitant with the inhibition of autophagosome/lysosome fusion. Cells with depleted levels *ITGA5* or *TSPAN4* exhibited lower growth rates and more dead cells after exposure to CQ or AS than control cells (Fig. 5G,H). As cell death is not directly associated with the knockdown of *ITGA5* or *TSPAN4*, we hypothesized that these genes are involved in ER stress-induced cell death via R&M formation. Thus, we identified the cellular states when cells were exposed to stress stimuli, in this case CQ or AS treatment. We confirmed ER stress markers in CQ- or AS-treated *ITGA5* or *TSPAN4* knockdown cells. We found that cells could not alleviate ER stress in the case of *ITGA5* or *TSPAN4* knockdown, exhibiting increased levels of spliced *XBP1* mRNA (Fig. 5I). Taken together, we revealed that decreased R&M formation did not relieve ER stress in GBM cells exposed to CQ or AS (Fig. 5J).

**Fig. 5.**
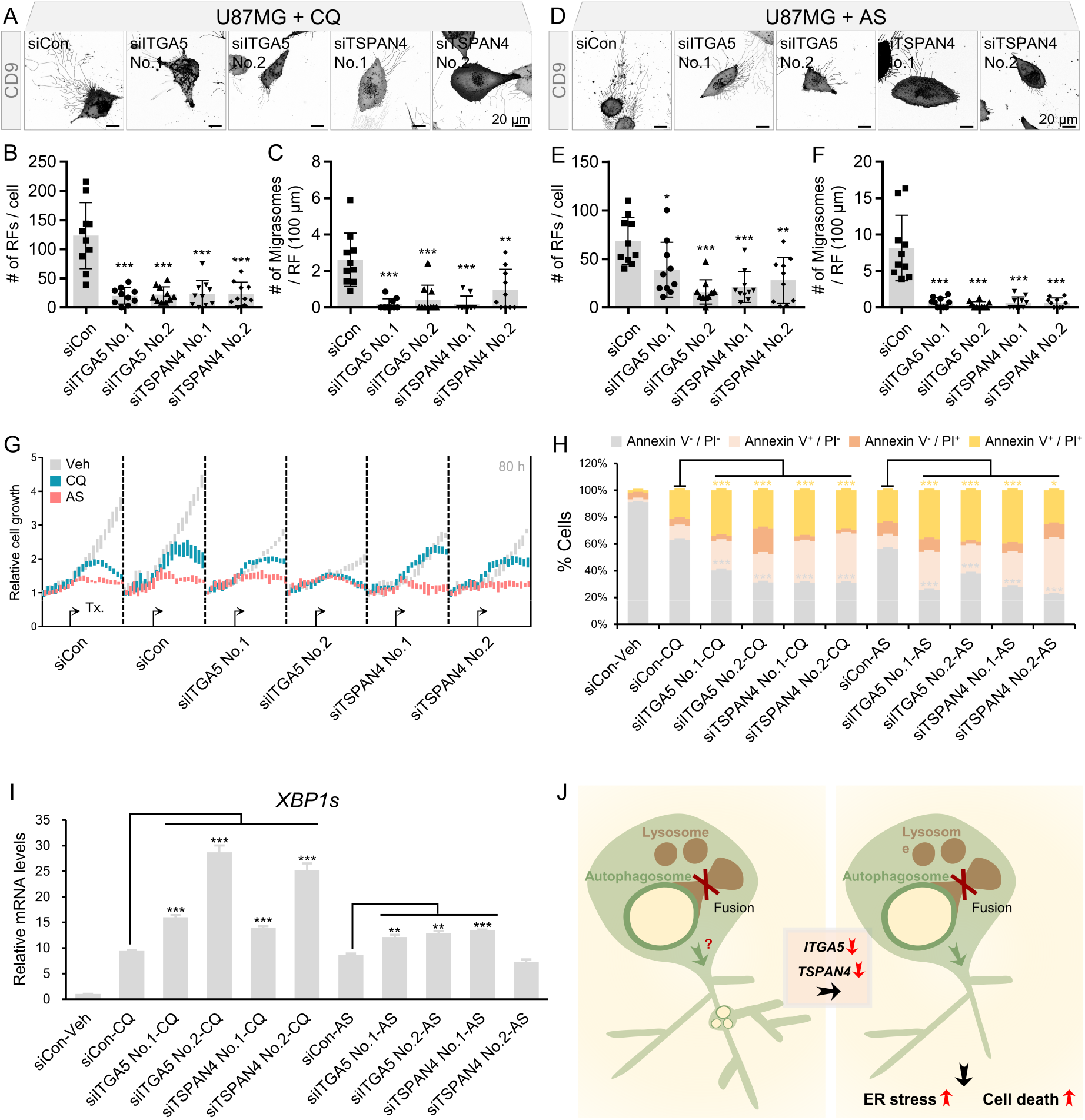
Cells restrained retraction fiber & migrasome (R&M) formation cannot alleviate ER stress. (A) Live-cell imaging of *ITGA5* or *TSPAN4* siRNA-transfected U87MG cells. Cells were treated with chloroquine (CQ; 50 μM, 12 h) or NaAsO_2_ (AS; 10 μM, 12 h). Scale bars, 20 μm. (B) Quantification of the number of RFs per a cell. Image analyses were performed using results from Fig. 5A. *** indicates p < 0.001. Data are expressed as mean ± SEM. The unpaired nonparametric Mann-Whitney U test was used to analyze the statistical significance between each group (n = 10). (C) Quantification of the number of migrasomes per RF (100 μm). Image analyses were performed using results from Fig. 5A. ** indicates p < 0.01. *** indicates p < 0.001. Data are expressed as mean ± SEM. The unpaired nonparametric Mann-Whitney U test was used to analyze the statistical significance between each group (n = 10). (D) Live-cell imaging of *ITGA5* or *TSPAN4* siRNA-transfected U87MG cells. Cells were treated with NaAsO_2_ (AS; 10 μM, 12 h). Scale bars, 20 μm. (E) Quantification of the number of RFs per a cell. Image analyses were performed using results from Fig. 5D. * indicates p < 0.05. ** indicates p < 0.01. *** indicates p < 0.001. Data are expressed as mean ± SEM. The unpaired nonparametric Mann-Whitney U test was used to analyze the statistical significance between each group (n = 10). (F) Quantification of the number of migrasomes per RF (100 μm). Image analyses were performed using results from Fig. 5D. *** indicates p < 0.001. Data are expressed as mean ± SEM. The unpaired nonparametric Mann-Whitney U test was used to analyze the statistical significance between each group (n = 10). (G) Cell growth rate of *ITGA5*- or *TSPAN4*-depleted U87MG cells in CQ (50 μM) or AS (10 μM)-treated conditions. Relative cell growth was calculated based on cell confluence mask analyzed by using IncuCyte ZOOM. (H) Annexin V/PI staining for analyzing cell death of *ITGA5*- or *TSPAN4*-depleted U87MG cells in CQ or AS-treated conditions. Cells were harvested after 72 h treatment of CQ or AS. * indicates p < 0.05; *** indicates p < 0.001. Data are expressed as mean ± SEM. Student’s t-test was used to analyze the statistical significance between each population (n = 3 for technical replicates). (I) qRT-PCR data for evaluating mRNA levels of spliced *XBP1*. * indicates p < 0.05. ** indicates p < 0.01. *** indicates p < 0.001. Data are expressed as mean ± SEM. Student’s t-test was used to analyze the statistical significance between each group (n = 3). (J) Graphical summary of study. In *ITGA5*- or *TSPAN4*-depleted condition, cells have a decreased number of R&Ms and lower capability to relieve ER stress, resulting in increased apoptosis.

### R&M are transferred to surrounding recipient cells and further degraded by lysosomes

We found that R&M are membranous structures produced under stressful conditions to mitigate cellular damage. In addition, we observed that these cellular structures could be transferred to neighboring cells. Thus, we further investigated the capacity of cells to take up R&Ms. There are many cell populations, including endothelial, neuronal, microglial cells, and mural cells (da Cunha et al., 2019). Therefore, we selected the following representative cell populations present in GBM lesions: astrocytes, endothelial cells, microglial cells, pericytes, and tumor cells. Owing to the adherent nature of R&Ms, we designed an R&M uptake assay in a co-culture system (Fig. S5A). In the co-culture assay, we analyzed the ratio of fragmented R&Ms on the bottom surface of culture dishes and the ratio of engulfed R&Ms inside the vesicles of recipient cells. We identified that microglial cells have a strong capacity to engulf R&Ms (Fig. S5B-E).

Next, we determined the exact endocytosis mechanism involved in R&M uptake by microglial cells. Microglial cells constitute a large proportion of the cellular mass within GBM lesions (∼30%) (Quail and Joyce, 2017). They also have diverse mechanisms of engulfing peripheral substances. Owing to their phagocytic ability, they usually take up many pathogens or other cellular materials via phagocytosis (Solé-Domènech et al., 2016). However, other endocytosis mechanisms, such as receptor-mediated endocytosis and macropinocytosis, are also activated in microglial cells (Solé-Domènech et al., 2016). Thus, we confirmed the modes of endocytosis involved in R&M uptake by microglial cells. Purified R&M labeled with PKH26 dye was used to track R&M uptake. PKH26-labeled purified R&Ms were gradually engulfed by BV2 microglial cells (Fig. S5F,G). R&M fluorescence was visualized 3 h after treatment. We then examined whether R&M uptake was inhibited by treatment with endocytosis inhibitors. Each endocytosis inhibitor, dynasore, ethylisopropyl amiloride (EIPA), and cytochalasin D, inhibited different modes of endocytosis. Among them, the dynamin GTPase inhibitor dynasore suppressed the uptake of R&M more efficiently compared to other inhibitors (Fig. S5H,I). The R&M uptake rate was gradually inhibited by increasing the dose of dynasore, indicating that receptor-mediated endocytosis is the dominant mechanism that serves the uptake capacity of BV2 microglial cells (Fig. S5J,K).

We further investigated the destination of engulfed R&M within the microglial cells. Thus, we treated BV2 microglial cells with purified R&M and performed immunofluorescence imaging to determine the destination of the engulfed R&Ms. We observed that R&Ms were localized with LAMP-1 positive lysosomes, not ER (calnexin-positive), suggesting that R&Ms are sent to lysosomes for further degradation (Fig. 6A). We also co-cultured LN229 cells expressing tandem-fluorescent LC3B with BV2 microglial cells to prove that LC3B-positive migrasomes of donor cells were engulfed and sequentially degraded by lysosomes of recipient cells. As expected, dual fluorescent-positive LC3B quenched the GFP signal in perinuclear lysosomes within BV2 microglial cells (Fig. 6B). In summary, we identified that microglial cells engulf R&M via receptor-mediated endocytosis and trafficked R&M toward lysosomes for further degradation (Fig. 6C).

**Fig. 6.**
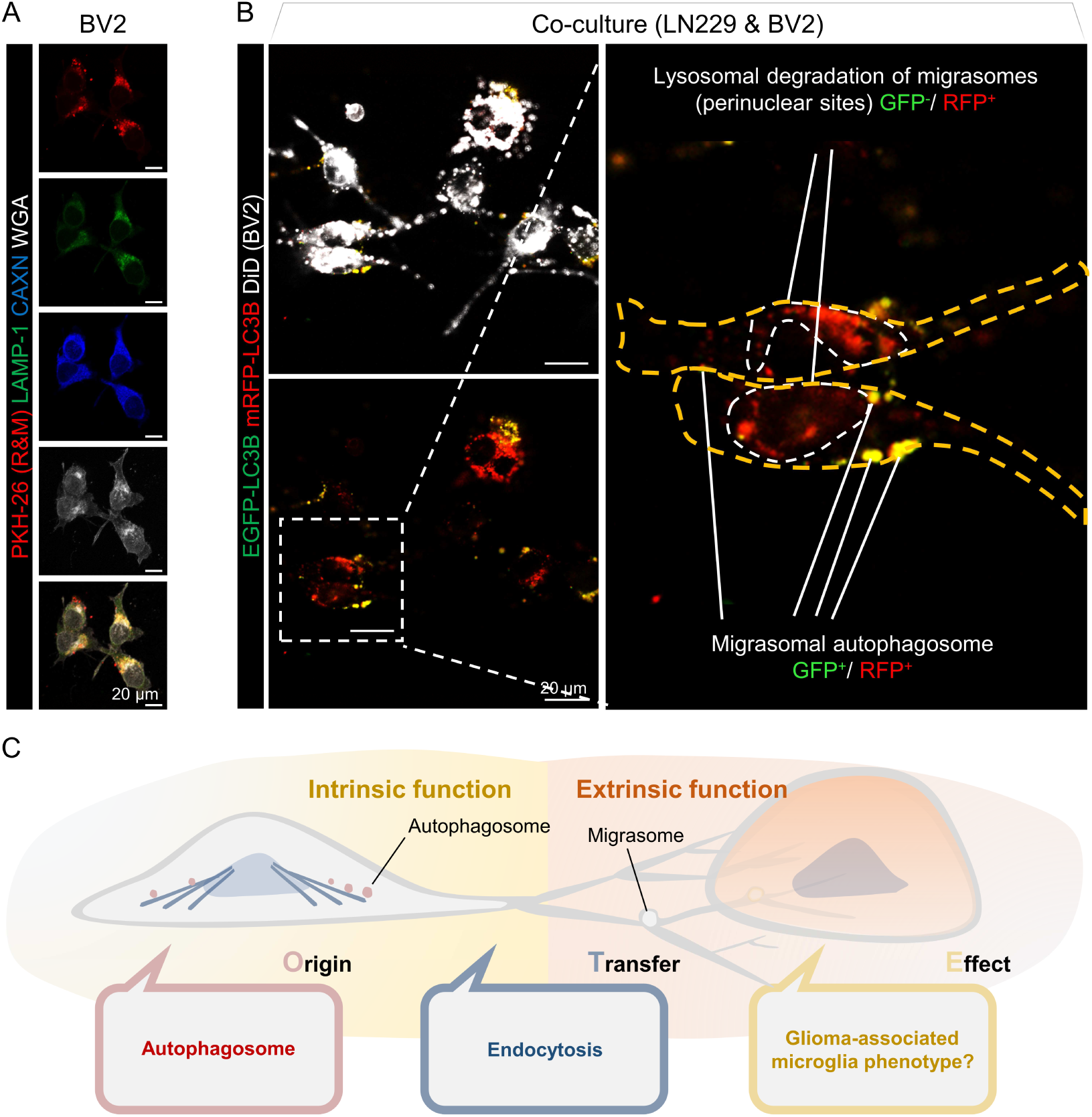
Engulfed retraction fibers & migrasomes (R&Ms) are destined to the lysosomes of recipient microglial cells. (A) Fluorescence imaging to detect the cellular location of purified R&M engulfed by BV2 microglial cells. Purified R&Ms were labeled by PKH26 dye. Red, PKH26-labeled purified R&M. Green, LAMP-1. Blue, Calnexin (CAXN). White, Wheat germ agglutinin (WGA). Scale bars, 20 μm. (B) Live-cell co-culture imaging for visualizing LN229 cells expressing tandem-fluorescent LC3B cultured with DiD-labeled BV2 microglial cells. Before co-culturing, LN229 cells were treated with chloroquine (CQ; 50 μM, 12 h). The culture medium was washed out before seeding BV2 cells. Cells were co-cultured for 12 h. Cells were imaged by z-stack live-cell confocal microscopy. White, DiD-labeled BV2 microglial cell. Green, EGFP-LC3B. Red, mRFP-LC3B. Scale bars, 20 μm. (C) Graphical summary of the study. Glioblastoma cell-derived tumor migrasomes contain autophagosomes, not autolysosomes. These cargo-containing vesicles can be transferred to surrounding microglial cells and eventually degraded by lysosomes.

## Discussion

In vivo environments always contain a matrix to which cells can attach (da Cunha et al., 2019; Sachs et al., 2017). Therefore, in such an environment, the formation of R&M occurs actively. In particular, tumor cells have a high potential to form cell membrane structures through cell migration or invasion. Our study revealed that the R&M formed by GBM cells was generated on the trailing side of the migrating cells and contained autophagosomes.

RFs appears to be a residue of actin filaments that the migrating cells have not retrieved. Actin filaments are known to interact with various proteins, and it was found that Arp2, one of the proteins constituting the branch of actin filaments, interacts with Atg9, a major factor involved in phagophore elongation (Monastyrska et al., 2008). The RF region, where the migrasome is present, is a junction point and seems to play a similar role to an actin filament branch. It is not known why these branch points are generated, but if it is assumed that the cytoskeleton is left behind by a migrating cell, the branch point will be a part of the Arp2/3 complex. The formation and maturation of autophagosomes cannot be considered in isolation, because of their association with the cytoskeleton (Kast and Dominguez, 2017). It has been reported that LC3 helps the actin nucleation function of JMY by binding to the LC3-interacting region (LIR) of the JMY protein during starvation (Hu and Mullins, 2019). Additionally, during the autophagy initiation phase, WHAMM, known as nucleation-promoting factor (NRF), recruits the Arp2/3 complex and is involved in autophagosome formation through the assembly of actin filaments aside from the ER (Kast and Dominguez, 2015). That is, the interaction between autophagosomes and the actin cytoskeleton is an essential process in the formation of autophagosomes, and the formation of migrasomes, which seem to have autophagosomes inherent in the RF passage, which is an actin-based structure, is a natural process.

In contrast, autophagosomes can fuse with vesicles constituting other endomembrane systems. A representative example is the fusion with endosome, which is called amphisome. The amphisome is LC3-positive and harbors endosomal markers (Jeppesen et al., 2019). The autophagosome bound to the multivesicular body shows a propensity to co-localize with exosomal markers. This type of fusion occurs in a unique environment such as shear stress (Wang et al., 2019). Amphisomes have macromolecules such as DNA and histones, separate from other EVs, and can be considered the origin of extracellular dsDNA (Jeppesen et al., 2019). Whether the LC3-positive vesicle found in the migrasome is an autophagosome or a different type of LC3-positive vesicle structure requires more precise identification.

The formation of autophagosomes and autophagy pathway are important mechanisms for alleviating the accumulation of stress damage in cancer cells (Kroemer et al., 2010; Ogata et al., 2006). Autophagosomes present in migrasomes may not function independently. However, it will help in the progression of the autophagy pathway, which appears through an increase in the spatial margin for autophagy, that is, the total cell area. In addition, instead of processing the accumulated autophagosome using the energy of the cell itself, the method of perishing from the cell body through the migrasome generated from cell movement is considered to support the survival of cancer cells.

A migrasome containing autophagosomes formed from cancer cells has the potential to serve as a nutrient source for recipient cells. In our study, microglial cells were the main recipient cells of R&M, suggesting the feasibility of affecting the polarity of microglial cells. Through our research, we confirmed that a number of ribosomes have been synthesized and have fulfilled their function for the vigorous translational needs of cancer cells in migrasomes. Therefore, if ribosomes composed of mega-complexes are delivered to microglial cells, it can be deduced that they provide energy sources or building blocks for microglial cells. Ribosomes are composed of a high ratio of amino acids, such as arginine and lysine (Wyant et al., 2018). In microglial cells, arginine is metabolized by NOS (nitric oxide synthase) or ARG1 (arginase-1), which are major proteins that regulate the characteristics of the known M1 and M2 types of microglial cells (Rath et al., 2014). Thus, it can be inferred that the migrasome, which is used as a raw material for providing amino acids, affects microglial polarity under certain circumstances.

Obviously, cells that make up any tissue do not exist statically. In this system, cells show dynamic movement, and in this process, more or less traces will be left, depending on the physical factors of the surrounding environment. Perhaps the cells were in constant communication through their dynamic movements and the substances they left behind, RF and migrasome. In the tumor environment, this membranous material exchange creates a favorable environment for cancer cells.

## Materials and Methods

### Cell lines and cell culture

Human GBM cell lines U87MG (male, RRID: CVCL_0022), LN229 (female, RRID: CVCL_0393) were purchased from the American Type Culture Collection (ATCC, Manassas, VA, USA). Murine brain microglial cell line BV-2 (female, RRID: CVCL_0182) was gifted from Dr. Jong Bae Park (National Cancer Center, Korea). All cell lines were authenticated using short tandem repeat (STR) profiling. The cells were cultured in high-glucose Dulbecco’s Modified Eagle medium (DMEM; Lonza, Basel, SWZ) supplemented with 10% fetal bovine serum (FBS; Hyclone, Logan, USA), 1% penicillin/streptomycin (P/S; Hyclone), and 2 mM L-glutamine (Hyclone) at 37 °C with 5% CO_2_ and 95% humidity.

### PLA

PLA was performed to detect LC3B-ATG5 binding in GBM cells. U87MG and LN229 cells (3 × 10^4^) expressing EGFP-CD9 were seeded on a coverslip in 48-well plates for 1 day. The cells were then fixed in 4% PFA for 15 min at RT. Cells were washed twice with 1x PBS. The fixed cells were permeabilized using 0.05% saponin (Sigma-Aldrich, Cat. # S7900, St. Louis, MO, USA) in PBS for 30 min. Samples were stained using primary antibodies for 3 h at RT followed by Duolink in situ PLA probes with anti-rabbit MINUS (Sigma-Aldrich, Cat. # DUO92005), anti-mouse PLUS (Sigma-Aldrich, Cat. # DUO92001), and Duolink in situ detection reagents Red (Sigma-Aldrich, Cat. # DUO92008). Anti-LC3B antibody (1:100; Novus Biologicals, Cat. # NB100-2220, Centennial, CO, USA) and anti-ATG5 antibody (1:100; Santa cruz Biotechnology, Cat. # sc-133158, Dallas, TX, USA) were used as primary antibodies. Mounted samples were observed using confocal laser scanning microscopy (CLSM) (LSM700; Carl Zeiss, Jena, DEU, Plan-Apochromat ×63/1.40 Oil DIC M27).

### Plasmids and lentivirus infection

EGFP-CD9, tdTomato-RPS13, tdTomato-RPLP2, were cloned into the pCDH-CMV-MCS-EF1a-Puro vector for overexpression. mRFP-EGFP-LC3B was cloned into the pLL-CMV-Puro vector for overexpression. sgRNA targeting *ATG5* (target sequence: No.1, 5′-AAGAGTAAGTTATTTGACGT-3′; No.2, 5′-CCTTAGATGGACAGTGCAGA-3′; No.3, 5′-TGATATAGCGTGAAACAAGT-3′), *ATG7* (target sequence: No.1, 5′-TCCTACTTTAGACTTGGACA-3′; No.2, 5′-CTTGAAAGACTCGAGTGTGT-3′; No.3, 5′-AAAGCAACAACATACCACAC-3′), and *BECN1*(target sequence: No.1, 5′-GGAAGAGACTAACTCAGGAG-3′; No.2, 5′-GAACCTCAGCCGAAGACTGA-3′; No.3, 5′-AGTGGCAGAAAATCTCGAGA-3′) were cloned into the pLentiCRISPRv2-Puro vector. Non-targeting sgRNA (5′-GCCCCGCCGCCCTCCCCTCC-3′) cloned into the pLentiCRISPRv2-Puro vector was used as a control. For transient expression, the plasmids were transfected using LipoJet (SignaGen Laboratories, Frederick, MD, USA). To construct a stable cell line, we performed lentivirus infection. To produce the lentivirus, each expression vector was transfected into HEK293T cells with second-generation lentiviral packaging plasmids pdR8.91 and pVSV-G using LipoJet. 24 h after transfection, the culture medium was harvested, incubated with Lenti-X concentrator (Clontech Laboratories, Cat. # h631231, Mountain View, CA, USA), and centrifuged to obtain the concentrated lentivirus. The cells were infected with the lentiviruses in the presence of 6 μg/mL polybrene (Sigma-Aldrich, Cat. # H9268) for 24 h.

### siRNA transfection

To perform siRNA-mediated knockdown of integrin genes, we transfected commercially available siRNAs (Sigma-Aldrich) using ScreenFect A Transfection Reagent (Wako Pure Chemical Industries, Cat. # 293-73201, Osaka, JPN) according to the manufacturer’s instructions. All siRNAs were synthesized from BIONEER (Double Strand RNA Oligo; BIONEER Inc., Daejeon, KOR). The siRNA information if as follows: human *STX17*, 5′-GAACACAAGUAUAUCAAGA-3′ and 5′-CUGGAAAUGUGAAAACUGA-3′; human *SNAP29*, 5′-GCAAAAUGCUUAUUAGAGU-3′ and 5′-GAUUUCCACUCUAUUGUGA-3′; human *ITGA5*, 5′-CUCCUAUAUGUGACCAGAG-3′ and 5′-GUUUCACAGUGGAACUUCA-3′; human *TSPAN4*, 5′-CCAUCGCCAUCCUCUUCUU-3′ and 5′-GUGGACCCCUCACCUACAU-3′.

### Electron microscopy imaging

Scanning electron microscopy was performed as previously described (Kim et al., 2012). Briefly, purified U87MG R&Ms were fixed using 2.5% glutaraldehyde (Sigma-Aldrich, Cat. # 354400) in 0.1 M phosphate buffer (pH 7.4) overnight at 4 °C. U87MG-derived brain tumor-bearing mice were anesthetized and perfused by the same fixative buffer. The samples were post-fixed with 2% osmium tetroxide for 2 h. After standard dehydration in ethanol series (60%, 70%, 80%, 90%, 95%, 100%, and 100%, each for 20 min, and then 100% for 30 min), the samples were immersed in t-butyl alcohol for 20 min twice. Next, the samples were dried using a freeze dryer (Hitachi ES-2030; Hitachi, Tokyo, JPN), and coated with platinum in ion sputter (Hitachi E-1045; Hitachi). The cells were observed using a scanning electron microscope (Hitachi S-4700; Hitachi).

For transmission electron microscopy, U87MG and LN229 cells were cultured on coverslips coated by FN. Next day, the experiment was performed as previously described (Kim et al., 2019). Briefly, the cells were fixed by 2% paraformaldehyde (PFA) /2.5% glutaraldehyde in 0.1 M phosphate buffer (pH 7.4) overnight at 4 °C. The samples were post-fixed with 2% osmium tetroxide for 2 h. After standard dehydration in ethanol series (60%, 70%, 80%, 90%, 95%, 100%, and 100% each for 20 min, and 100% for 30 min), the dehydrated samples were suspended in propylene oxide for 20 min. Epon infiltration was performed by the serial incubation of 2:1, 1:1, 1:2 of propylene oxide:epon mixture for 1 h each. Next, the samples were embedded in epoxy resin mixture and polymerized at 60 °C in a dry oven for 48 h. Polymerized blocks were fine trimmed into the paramedian lobule. Semi-thin sections (1 µm) were obtained using an ultramicrotome (Leica UC7; Leica, Wetzlar, DEU), and observed via toluidine blue staining. Sections were mounted on Formvar-coated one-hole grids and subsequently post-stained with heavy metals. Serial images were randomly obtained from the samples using a transmission electron microscope (Hitachi H-7650; Hitachi).

### Gene Ontology (GO) analysis

R&M-specific Q1 gene sets from LC-HRMS data were analyzed using the ClueGO v2.5.4 plugin (Bindea et al., 2009). This software was plugged into the CytoScape v3.6.1 (Shannon et al., 2003). To group GO terms, the kappa score was set at 0.4 and the number of overlapping genes to combine groups was set at 50%.

### Gene interactive network analysis

In silico prediction of the possible genes association network and genes functions was performed using the GeneMANIA v3.5.1 (Gene Function Prediction using a Multiple Association Network Intergration Algorithm) (Warde-Farley et al., 2010).

### Live-cell fluorescence imaging and quantification of R&Ms

GBM cells (3 × 10^4^) were seeded on a coverslip in 24-well plates for 1 day. The cells were then imaged using live-cell confocal microscopy. Live cell images were obtained using CLSM (LSM700; Carl Zeiss, Plan-Apochromat ×63/1.40 Oil DIC M27, ×10-35 images of U87MG cells and ×10-35 images of LN229 cells were obtained by 1× magnification at ×63, ZEN acquisition software version 2018, blue edition) with live cell incubator (Chamlide TC-FC5N; Live Cell Instrument (LCI), Seoul, KOR). he images were randomly taken for each sample. We used ImageJ software v.1.52a (National Institute of Health (NIH), Bethesda, MD, USA) to calculate the RFs, their junctions, total length, and the number of migrasomes of all images by using “Ridge Detection” plugin (Steger, 1998). CQ (Selleckchem, Cat. # NSC-187208, Houston, TX, USA), AS (Alfa Aesar, Cat. # 041533.AP, 0.1N standardized solution, Ward Hill, MA, USA), and BafA1 (Sigma-Aldrich, Cat. # B1793) were used for experiments.

### Immunofluorescence

BV2 cells were seeded on a coverslip in 24-well plates for 1 day. The cells were then fixed in 4% PFA and incubated with wheat germ agglutinin (WGA; 5.0 μg/mL) for 15 min at RT. Cells were washed twice with 1× PBS. For immunofluorescence staining for LAMP-1 and calnexin, the fixed cells were permeabilized by 0.1% Triton X-100 in PBS for 15 min at RT, then blocked by 5% BSA in PBS for 1 h at RT. Cells were incubated with LAMP-1 antibody (1:200; abcam, Cat. # ab25630, Cambridge, MA, USA) and calnexin antibody (1:200; GeneTex, Cat. # GTX109669, Irvine, CA, USA) overnight at 4 ℃, followed by incubating Alexa Fluor 350 or 488-conjugated secondary antibody (1:500) for 2 h at RT. Cell images were obtained using CLSM (LSM700).

### Sample preparation for proteomics

EV and R&M samples were resuspended in 400 μL of 5% SDS in 50 mM TEAB (pH 7.55), and dithiothreitol was added to a final concentration of 20 mM for 10 min at 95 °C to reduce disulfide bonds. Reduced samples were then incubated with 40 mM iodoacetamide for 30 min at room temperature in the dark. By a 10-fold dilution of 12% phosphoric acid, acidified samples were loaded onto S-Trap macro (EV) or mini (R&M) columns (ProtiFi, Farmingdale, NY, USA; Cat. No: CO2-macro-80 or CO2-mini-80). We treated suspension-trapping (S-trap) proteolysis according to the manufacturer’s protocol, followed by the addition of 1:20 Lys-C/trypsin mixture and incubation for 16 h at 37 °C (HaileMariam et al., 2018). The eluted peptide mixture was lyophilized using a cold trap and stored at -80 °C until use.

### Nano-LC-ESI-MS/MS analysis

The LC system was an Dionex UltiMate 3000 RSLCnano system (Thermo Fisher Scientific). Mobile phase A was 0.1% formic acid and 5% DMSO in water and mobile phase B was 0.1% formic acid, 5% DMSO and 80% acetonitrile in water. Samples were reconstituted with 25 µL of mobile phase A, injected with a full sample loop injection of 5 µL into a C18 Pepmap trap column (20 × 100 μm i.d., 5 μm, 100 Å; Thermo Fisher Scientific), and separated in EASY-Spray column (500 × 75 μm i.d., 2 μm, 100 Å; Thermo Fisher Scientific) over 200 min (250 nL/min) at 50 °C. The column was priory equilibrated with 95% mobile phase A and 5% mobile phase B. A gradient of 5–40% B for 150 min, 40–95% for 2 min, 95% for 23 min, 95–5% B for 10 min, and 5% B for 15 min were applied. The LC system was coupled to an Orbitrap Exploris 480 mass spectrometer (Thermo Fisher Scientific) with a nano EASY-Spray™ source. For all experiments, spray voltage was set to 2.1 kV, ion transfer tube temperature to 290 °C, and a radio frequency funnel level to 40%. Full scans were made at a resolution of 60,000. MS1 AGC target was set to 3,000,000 charges, with a maximum injection time of 25 ms. Scan ranges used (in m/z) were as follows; 350–1600. Precursor fragmentation was achieved through higher-energy collision disassociation (HCD) with a normalized collision energy of 30, and an isolation width of 1.3 m/z. Full scans were made at a resolution of 15,000. MS2 AGC target was set to 2,000,000. Charges 2–5 were considered for MS/MS, with a dynamic exclusion of 20 ms. Loop counts were set to 12. MS2 resolution was set to 15,000. Maximum MS2 injection times were as follows: 22 ms.

### Protein Identification by database search

Individual raw files acquired MS analysis and were retrieved against the reviewed Human Uniprot-SwissProt protein database (released on June 2019) (2021) using the SEQUEST-HT on Proteome Discoverer (Version 2.2, Thermo Fisher Scientific). Search parameters used were as follows: 10-ppm tolerance for precursor ion mass and 0.02 Da for fragmentation mass. Trypsin peptides tolerate up to two false cleavages. Carbamidomethylation of cysteines was set as fixed modification and N-terminal acetylation and methionine oxidation were set as variable modifications. The false discovery rate was calculated using the target-decoy search strategy, and the peptides within 1% of the FDR were selected using the post-processing semi-supervised learning tool Percolator (Käll et al., 2007) based on the SEQUEST result. Label-free quantitation of proteins was calculated using the precursor ion peak intensity for unique and razor peptides of each protein and excluded peptides with methionine oxidation.

### Western blotting

Western blot was performed to analyze the protein expression. Briefly, cell extracts were prepared using RIPA lysis buffer (150 mM sodium chloride, 1% NP-40, 0.1% SDS, 50 mM Tris, pH 7.4) containing 1 mM β-glycerophosphate, 2.5 mM sodium pyrophosphate, 1 mM sodium fluoride, 1 mM sodium orthovanadate, and protease inhibitor (Roche, Cat # 11836170001, Basel, Switzerland). The protein concentration was quantified using the Bradford assay reagent (Bio-Rad, Hercules, CA, USA) according to the manufacturer’s instructions. Proteins were resolved by SDS-PAGE and then transferred to a polyvinylidene fluoride membrane (Pall Corporation, Port Washington, NY, USA). Membranes were blocked with 5% non-fat milk and incubated with the primary antibody. Membranes were then incubated with horseradish peroxidase-conjugated anti-IgG secondary antibody (Pierce Biotechnology, Rockford, IL, USA) and visualized using the SuperSignal West Pico Chemiluminescent Substrate (Pierce Biotechnology). Primary antibodies used for western blot analysis are as follows: anti-SQSTM1 (p62; 1:1000; Santa cruz Biotechnology, Cat. # sc-28359), anti-β-actin (1:10000; Santa cruz Biotechnology, Cat. # sc-47778), anti-Beclin-1 (1:1000; Santa cruz Biotechnology, Cat. # sc-48341), anti-ATG7 (1:500; Santa cruz Biotechnology, Cat. # sc-376212), anti-ATG5 (1:500; Santa cruz Biotechnology, Cat. # sc-133158), anti-LC3B (Novus Biologicals, Cat. # NB100-2220), anti-GRP78 (BiP; 1:1000; abcam, Cat. # ab212054), anti-HSC70 (1:1000; Santa cruz Biotechnology, Cat. # sc-7298), anti-RPS13 (1:500; Santa cruz Biotechnology, Cat. # sc-398690), anti-RPL4 (1:500; Santa cruz Biotechnology, Cat. # sc-100838), anti-α-tubulin (1:10000; Sigma-Aldrich, Cat. # T6199), anti-integrin α5 (1:1000; abcam, Cat. # ab150361), anti-CD63 (1:1000; Santa cruz Biotechnology, Cat. # sc-15363), anti-p-eIF2α (Ser51) (1:1000; Cell signaling, Cat. # 9721, Danvers, MA, USA).

### Quantitative reverse transcription-PCR (qRT-PCR)

The qRT-PCR was performed to determine mRNA levels. Briefly, total RNA was isolated from cells using the QIAzol lysis reagent (QIAGEN, Cat. # 79306, Valencia, CA, USA) according to the manufacturer’s instructions. The 1 U of DNase I, RNase-free (Thermo Fisher Scientific, Cat. # EN0525), was added to 1 μg of template RNA, and incubated for 30 min at 37 ℃. For inactivating DNase I, 50 mM of EDTA was treated and heated at 65 ℃ for 10 min. DNase I treated RNA was utilized as a template for synthesizing complementary DNA (cDNA) using the RevertAid First Strand cDNA Synthesis Kit (Thermo Fisher Scientific, Cat. # K1622) according to the manufacturer’s instructions. The qRT-PCR analysis was performed using the Takara Bio SYBR Premix Ex Taq (Takara, Cat. # RR420A, JPN) and CFX096 (Bio-Rad). The expression levels of each target gene were normalized to that of 18S rRNA. The primers used for the analyses are listed in Supplementary Table 1.

### Cell death assay

Cell death was evaluated using flow cytometry analysis. Briefly, U87MG (2.5 × 10^5^) cells were seeded in a 60-mm culture dish and incubated with CQ or AS for 72 h. Cell pellets were centrifuged at 3500 rpm for 5 min and then stained with APC Annexin V (BD Biosciences Pharmingen, Cat. #550474, Franklin Lakes, NJ, USA). Each cell suspension (100 μL in binding buffer) stained with 10 μL APC Annexin V and 20 μL propidium iodide (PI; 50 μg/mL) was gently mixed and incubated for 15 min. Next, an additional 200 μL of binding buffer was added, then the cells were immediately analyzed using a FACSVerse apparatus. Non-apoptotic cell death (cells stained with PI alone), early apoptosis (cells positive for APC AnnexinV alone), and late apoptosis (cells positive for PI and APC Annexin V) were observed.

### Statistical analysis

All data from the experiments, shown as bar graphs, are presented as mean ± SEM. Statistical analysis was performed by using the unpaired nonparametric Mann-Whitney U test or two-tailed Student’s t-tests. Values of p < 0.05 or p < 0.01 were considered statistically significant for different experiments, as indicated in the figure legends.

## Acknowledgments

We are grateful to all members of the Cancer Growth Regulation Laboratory for their helpful discussion and technical assistance. We thank Minsu Jang for his kind provide of experimental material (ptfLC3 vector, tandem-fluorescent LC3B).

## Author Contributions

S. Y. Lee and H. Kim designed the experiments and wrote the manuscript. S. Y. Lee and S. -H. Choi conducted most experiments and analyzed the data. H. -S. Ahn and K. Kim performed proteomic analysis and relevant experiments. S. W. Chi established bioinformatic data and relevant experiments. Y. -G. Ko gave us motivation to investigate this study. H. Kim directed this study.

## Competing interests

No competing interests declared.

## Funding

This study was supported by grants to S. W. Chi and H. Kim from the Science Research Center Program through the National Research Foundation of Korea (NRF) funded by the Ministry of Science and ICT (MSIT) (NRF-2015R1A5A1009024), to K. Kim from NRF funded by MSIT (NRF-2019M3E5D3073106), to and to the School of Life Sciences and Biotechnology for BK21, Korea University.

## Notes

### Competing Interest Statement

The authors have declared no competing interest.

